# Fairness-aware Supervised Hierarchical Contrastive Semantic Learning for Sexual Dimorphism Analysis

**DOI:** 10.64898/2026.01.27.702125

**Authors:** Euiseong Ko, Sai Phani Parsa, Sai Chandra Kosaraju, Tesfaye B. Mersha, Mingon Kang

## Abstract

**Motivation:** Sexual dimorphism is a fundamental biological determinant driving systematic differences in disease susceptibility, progression, and clinical outcomes. However, current AI-based genomic models often exhibit algorithmic bias and fail to capture these sex-specific mechanisms, creating a critical barrier to unbiased precision medicine. Ensuring fairness in the context of sexual dimorphism requires understanding and addressing the distinct biological mechanisms functioning in each sex, rather than focusing solely on equalizing predictive performance.

**Results:** We propose a fairness-aware supervised hierarchical contrastive learning approach, called FairHICON, to discover unbiased sex-common and sex-specific genomic drivers. Evaluations on cancer and asthma transcriptomic datasets demonstrate that FairHICON significantly outperforms state-of-the-art benchmarks, improving predictive performance by up to 9% while effectively reducing the performance gap between male and female cohorts. Furthermore, prognostic validation confirms that the identified sex-specific pathways stratify patient survival significantly better within their corresponding sex groups. This validates FairHICON to elucidate the molecular heterogeneity of sexual dimorphism, advancing inclusive precision medicine.

**Availability and implementation:** The source code and data is available at https://github.com/datax-lab/FairHICON.

## 1. Introduction

Biological sex significantly influences human physiology, leading to substantial differences in disease susceptibility, progression, and clinical outcomes (Ober et al., 2008; Jun et al., 2021; Kim et al., 2018; Zheng et al., 2019). For instance, asthma prevalence shifts during development; it is more common in boys during childhood, but affects women more frequently and severely after puberty, indicating a specific interaction between sex hormones and immune function (Postma, 2007). In oncology, men generally show higher incidence rates and lower survival probabilities than women for various cancers, including those of the bladder, colon, liver, and brain (Rubin et al., 2020; Cook et al., 2011; Siegel et al., 2021). These phenotypic observations stem from distinct molecular mechanisms, characterized by divergent regulatory networks and pathogenic pathways that differ between males and females (Oliva et al., 2020; Lopes-Ramos et al., 2020, 2018; Hartman et al., 2021).

Analytical methodologies have advanced in elucidating the molecular mechanisms underlying sexual dimorphism. Conventional statistical approaches, such as differential expression analysis, established the foundation for identifying genes associated with sex-specific traits (Gautam et al., 2019; Bourquard et al., 2023). Mostly, these studies have relied on independent analyzes that examine male and female cohorts separately to isolate sex-specific associations. The field has shifted towards advanced deep learning models to capture non-linear relationships in high-dimensional genomic data (Hases et al., 2021; Ko et al., 2024). For instance, the Sex-specific and Pathway-based Interpretable Neural network (SPIN) incorporates biological pathway information to detect sex-specific molecular features in cancer and asthma (Ko et al., 2024). These advances demonstrate the increasing capacity of AI to characterize the genomic basis of sex differences.

Despite these methodological advances, the application of AI to translational genomics faces significant challenges in terms of algorithmic fairness. Most deep learning models are susceptible to reinforcing inherent biases from data, particularly when training data contain substantial imbalances between male and female populations (Ober et al., 2008). These models frequently exhibit skewed bias toward the majority group or rely on misleading statistical associations between sex and disease, rather than uncovering stable, causal molecular mechanisms. Such biases manifest as clear differences in performance, providing stronger predictive power for one sex while reducing how well the model generalizes to the other.

More importantly, fairness in sexual dimorphism focuses on reducing the knowledge gap as well as disparity in predictive performance across sex groups. Achieving statistical parity in predictions does not guarantee fairness in model interpretation, which is critical in the sexual dimorphism analysis. Identical clinical phenotypes frequently arise from distinct underlying molecular mechanisms in males and females (Oliva et al., 2020; Lopes-Ramos et al., 2020, 2018). Reducing the performance gap between the sex groups without accounting for these distinct drivers can obscure the distinct pathogenic mechanisms present in the minority group, thereby limiting the clinical relevance of the findings.

To address this critical gap, we propose a novel approach, Fairness-aware Supervised Hierarchical Contrastive Semantic Learning (FairHICON), to mitigate sex bias while uncovering the discovery of unbiased sex-common/-specific genomic mechanisms. FairHICON introduces a novel two-level hierarchical supervised contrastive loss designed to identify and characterize the specific molecular determinants present in each sex group. This objective methodically refines the structure of the latent space by using phenotypic labels to maintain clear separation between classes, while simultaneously exploiting sex attributes to enforce disentanglement across subgroups. By further integrating adaptive prototype weighting with pathway architectures informed by biological prior knowledge, FairHICON ensures that the learned representations remain both statistically fair and biologically interpretable for both sexes. The main contributions of FairHICON are (1) a novel fairness-aware approach integrating supervised hierarchical contrastive learning with biologically informed architectures to ensure unbiased representation learning; (2) superior predictive performance that effectively closes the accuracy gap between male and female cohorts compared to state-of-the-art benchmarks; and (3) fair and interpretable discovery, enabling the identification of clinically relevant sex-common and sex-specific biological risk factors for both sexes.

## 2. Materials and Methods

We introduce FairHICON that mitigates sex disparities in predictive performance and model interpretation by learning unbiased sex-specific and sex-common transcriptomic mechanisms. The following sections outline the theoretical basis of contrastive learning for the analysis of sexual dimorphism, describe the network architecture and the hierarchical objective function of FairHICON, and present the training strategy used to enhance both stability and interpretability.

### 2.1. Contrastive learning for sexual dimorphism

Contrastive learning is a representation learning paradigm designed to optimize the geometry of a latent feature space. The fundamental principle of contrastive learning is to pull semantically similar samples (positive pairs) closer together while pushing dissimilar samples (negative pairs) apart. This learning process organizes data vectors based on their intrinsic relationships, creating a structured embedding space where the spatial distance directly corresponds to semantic similarity.

In the context of sexual dimorphism analysis, the contrastive learning provides a powerful mechanism for feature disentanglement. A typical application could define similarity based on a straightforward pairing of clinical outcome and sex (e.g., treating male-disease and female-disease as completely separate categories). However, this combinatorial method has a significant drawback. By treating these subgroups as strictly negative pairs, the standard contrastive objective forces the model to separate them in the latent space. Although this rigorous separation highlights sex-specific distinctions, it inadvertently suppresses sex-common biological mechanisms that are fundamentally shared throughout the disease regardless of sex. As a result, a straightforward combinatorial method cannot simultaneously preserve these conserved signals and separate sex-specific heterogeneity at the same time, making it necessary to adopt a hierarchical strategy that can balance these opposing objectives.

### 2.2. The FairHICON architecture

FairHICON employs a multi-branch Siamese architecture designed to explicitly disentangle sex-specific mechanisms from shared disease drivers (Fig. 1). Given the labeled dataset, let 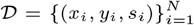, where *x*_*i*_ represents the gene expression profile (e.g. RNA-seq), *y*_*i*_ ∈ {0, 1} denotes the clinical outcome, and *s*_*i*_ ∈ {*M, F*} represents the sex attribute.

**Fig. 1.**
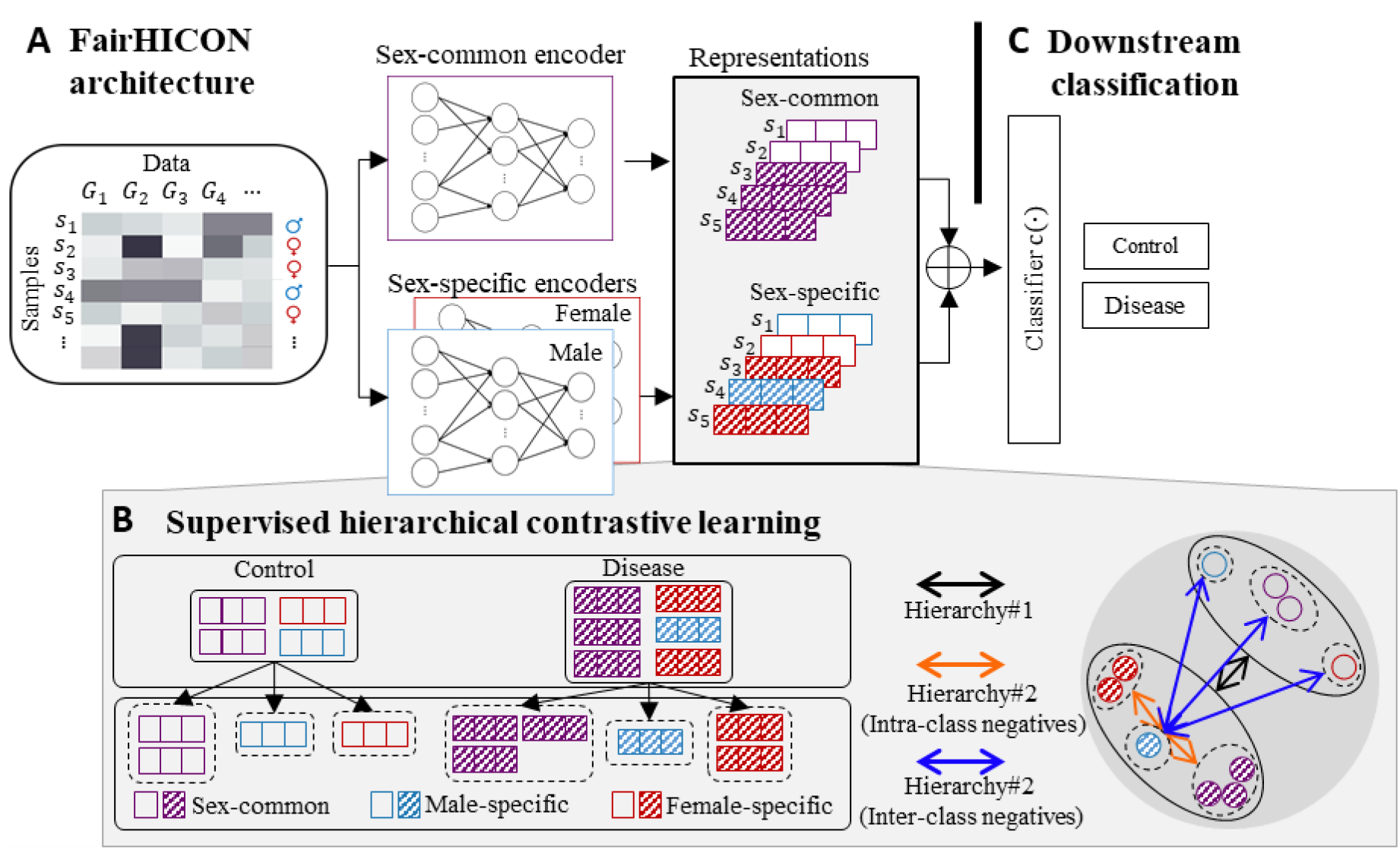
Overview of FairHICON. The approach takes gene expression data and sex attribute as input, processing them through a Siamese architecture. The sex-common backbone model extracts features shared across groups, while sex-specific models (male and female) capture sex-specific patterns. These representations are optimized via supervised hierarchical contrastive learning (bottom panel), which organizes the latent space representations into two levels: Level 1 Hierarchy (Hierarchy#1) separates samples based on the primary clinical outcome (control vs. disease), while Level 2 Hierarchy (Hierarchy#2) refines the feature space using hard negatives (blue arrows) to distinguish subgroup representations between different classes, and soft negatives (orange arrows) to disentangle subgroup features within same class. These representations are fused (⊕) to perform the downstream tasks, specifically cancer survival prediction (classifying Long-Term Survivors, LTS, vs. Non-LTS) and asthma risk score prediction (stratifying patients).

The architecture consists of three distinct encoders: a sex-common encoder (*E*_*C*_) to capture sex-common features, and two sex-specific encoders (*E*_*M*_ and *E*_*F*_) to capture mechanisms unique to males and females. To ensure biological interpretability, each encoder is adapted using the Pathway-Associated Sparse Deep Neural Network (PASNet) architecture (Hao et al., 2018). Unlike standard fully connected networks, PASNet enforces sparse connectivity constrained by prior biological knowledge (e.g., KEGG pathway annotations). This design ensures that the internal topology of the network mirrors hierarchical biological systems, allowing the direct attribution of predictive signals to specific molecular pathways. The information flows through three specific transformation stages:

The gene-to-pathway layer constitutes the input stage, where *d*_*g*_ nodes correspond to individual genes mapping to *d*_*p*_ nodes representing biological pathways. Following the PASNet structure, the weight matrix *W* ^(1)^ is constrained by a binary mask 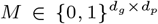, where *M*_*jk*_ = 1 if and only if gene *j* is annotated to pathway *k*. The forward propagation for the pathway activation vector *h* is computed as:

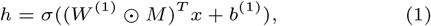

where, ⊙ denotes the element-wise Hadamard product, which strictly zeroes out weights corresponding to non-existent biological relationships.

Subsequently, the pathway-to-representation layer aggregates the pathway activations *h* into a lower-dimensional latent embedding vector *z*_*r*_ ∈ ℝ via a fully connected layer. This layer synthesizes the activity of disparate biological processes into a compact representation:

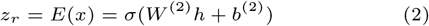

Finally, during the contrastive pre-training phase, a projection head *H*(·) maps the representation into the metric space where the loss is calculated. This head consists of a two-layer MLP (*z*_*proj*_ = *W* ^(4)^*σ*(*W* ^(3)^*z*_*r*_)). This design decouples representation learning from contrastive optimization, allowing the encoder to retain the information necessary for downstream tasks, while the projection head absorbs the geometric distortions induced by contrastive loss (Chen et al., 2020).

For every patient sample *i*, the approach generates a dual-view representation comprising a sex-common view, denoted as 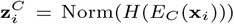, and a sex-specific view, denoted as 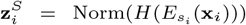. Here, *H*(·) is the projection head and Norm(·) denotes *L*_2_ normalization. These views are aggregated into an augmented batch *Z* of size 2*N*, indexed by *k* ∈ {1, …, 2*N*}. Each embedding in the batch is explicitly associated with a semantic triplet (*y*_*k*_, *s*_*k*_, *t*_*k*_), where *t*_*k*_ ∈ {common, male-/female-specific} denotes the encoder origin.

### 2.3. Supervised hierarchical contrastive semantic loss

We propose the Supervised Hierarchical Contrastive Semantic Loss (SHCSL) that learns discriminative representations of clinical outcomes and invariant to sex-based biases. This objective function is formulated in the two hierarchical levels of inter-class separability and intra-class disentanglement.

The total objective function aggregates the hierarchical level losses (*h*1 and *h*2) which are computed on both the projection space and the representation space. The total loss is defined as:

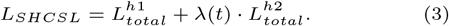

The hierarchical losses consider four semantic relationship between an anchor sample *u* and any other sample *v*, defined as:

- Intra-Class, Intra-Attribute (𝒮_*ICIA*_) is the similarity between an anchor *u* and samples belonging to the exact same subgroup, defined as 𝒮_*ICIA*_(*u*) = {*v* ∈ 𝒵 *\ u* | *y*_*v*_ = *y*_*u*_ ∧ *s*_*v*_ = *s*_*u*_ ∧ *t*_*v*_ = *t*_*u*_}.
- Intra-Class, Inter-Attribute (𝒮_*ICEA*_) is the similarity between an anchor *u* and samples sharing the clinical label but differing in subgroup (different sex or encoder origin), defined as 𝒮_*ICEA*_(*u*) = {*v* ∈ 𝒵 \ {*u*} | *y*_*v*_ = *y*_*u*_ ∧ (*s*_*v*_ ≠ *s*_*u*_ ∨ *t*_*v*_≠ *t*_*u*_)}.
- Inter-Class, Intra-Attribute (𝒮_*ECIA*_) is the similarity between an anchor *u* and samples with different clinical labels but sharing sensitive attributes, defined as 𝒮_*ECIA*_(*u*) = {*v* ∈ 𝒵 \ {*u*} | *y*_*v*_≠ *y*_*u*_ ∧ *s*_*v*_ = *s*_*u*_}.

Inter-Class, Inter-Attribute (𝒮_*ECEA*_) is the similarity between an anchor *u* and samples differing in both label and attributes, defined as 𝒮_*ECEA*_(*u*) = {*v* ∈ 𝒵 \ {*u*} | *y*_*v*_≠ *y*_*u*_ ∧ *s*_*v*_ *s*_*u*_}.

The first level hierarchy maximizes global predictive accuracy by clustering samples based strictly on the clinical outcome. The positive set (𝒫_ℋ1_) includes all samples sharing the same clinical label *y*, regardless of sex or encoder origin, defined as 𝒫_ℋ1_(*u*) = 𝒮_*ICIA*_(*u*) ∪ 𝒮_*ICEA*_(*u*). We employ standard uniform weighting (*w*_*u,a*_ = 1) to learn robust decision boundaries. To mitigate the impact of group imbalance, the loss employs group-wise normalization: the loss is calculated independently for each unique subgroup and then averaged, ensuring that minority groups contribute equitably to the gradient (Park et al., 2022):

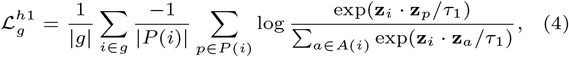

where *τ*_1_ is a standard temperature parameter.

The second level hierarchy promotes fairness by disentangling subgroups within the same disease class. The positive set (𝒫_ℋ2_) is restricted strictly to samples from the same semantic subgroup, such that 𝒫_ℋ2_(*u*) = 𝒮_*ICIA*_(*u*). To effectively structure the latent space, we categorize negative samples into two structural types with distinct optimization goals. Intra-class negatives (𝒩_*intra*_) are samples sharing the clinical label *y*_*u*_ but belonging to a different semantic subgroup (𝒩_*intra*_(*u*) = 𝒮_*ICEA*_(*u*)); these samples are structurally proximal due to ℋ_1_ but require active penalization to prevent the merging of sex-specific manifolds. Inter-class negatives (𝒩_*inter*_) are samples belonging to the opposing clinical class (𝒩_*inter*_(*u*) = 𝒮_*ECIA*_(*u*) ∪𝒮_*ECEA*_(*u*)), representing the fundamental decision boundary. To establish a stable reference for each semantic cluster, we compute a group prototype **c**_*g*_ for every unique subgroup *g* ∈ 𝒢. This prototype is defined as the *L*_2_-normalized centroid of all embeddings belonging to that subgroup:

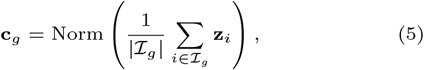

where ℐ_*g*_ denotes the set of indices for samples belonging to subgroup *g*, and Norm(·) applies *L*_2_ normalization.

We then employ an adaptive weighting mechanism, *w*_*u,a*_, to penalize negatives that violate the subgroup structure relative to the anchor’s group prototype **c**_*g*(*u*)_:

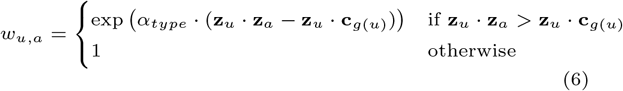

We assign distinct scaling factors, *α*_*intra*_ and *α*_*inter*_, to the respective negative sets. We set *α*_*intra*_ *< α*_*inter*_ to prioritize the stability of the global decision boundary (separating disease vs. control) while sufficiently penalizing intra-class overlap to enforce fine-grained, sex-specific disentanglement. To ensure numerical stability and adherence to column width constraints, we define the weighted denominator term *D*_*weighted*_(*i*) separately: The weighted contrastive loss is defined as:

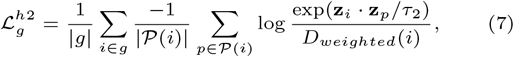

where the weighted denominator *D*_*weighted*_(*i*) aggregates contributions from positive pairs and weighted negative samples. Let *N* (*i*) be the set of negative indices for the anchor *i* (i.e., *N* (*i*) = *A*(*i*) *\ P* (*i*)). Then:

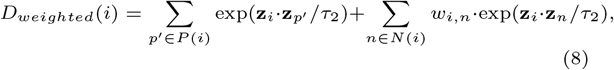

where *τ*_2_ is the temperature parameter for the second hierarchy.

To ensure the structural integrity of the learned features, the component losses for each hierarchy are calculated as a weighted sum of the loss on the projection and representation spaces, controlled by a balance parameter *α*:

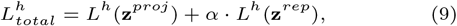

where the representations **z**^*rep*^ are *L*_2_-normalized specifically for this loss calculation.

### 2.4 Training Strategy

We employ a two-phase curriculum learning strategy to stabilize the optimization of the hierarchical objective (Wang et al., 2021). In Phase 1, the network was optimized exclusively using the first-level hierarchy loss 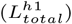 to establish robust disease classification boundaries between disease and control groups without the constraint of subgroup disentanglement. In Phase 2, the second-level hierarchy loss 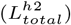 was introduced to enforce fairness and disentanglement. To prevent destabilization of the learned features, the weight *λ*(*t*) was linearly increased from zero to its target value over a defined warm-up period, after which it was held constant. This allows the model to gradually refine the latent space, separating sex-specific manifolds within the established class clusters. Model selection is performed via early stopping based on the silhouette score of the validation set representations, ensuring that the model converges to a state with optimal cluster integrity.

### 2.5. Downstream Binary Classification

Following the contrastive learning, the projection heads are discarded, and the BINN encoders (*E*_*C*_, *E*_*M*_, *E*_*F*_) are frozen to serve as feature extractors. For a patient *i* with sex *s*_*i*_, we generate a fused representation **v**_*i*_ by concatenating the unnormalized features from the sex-common and the corresponding sex-specific encoder:

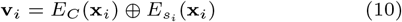

This joint vector **v**_*i*_ ∈ ℝ encapsulates both the common biological context and the specific characteristics of the sexual dimorphism. A classifier (e.g., a neural network) is subsequently trained on **v**_*i*_ to predict the binary clinical outcome *y*_*i*_. The classifier is optimized using standard Binary Cross-Entropy (BCE) loss.

### 2.6. Feature Attribution and Interpretation

To identify the specific genomic drivers of the model’s predictions, we leverage the interpretable BINN architecture. We compute importance scores for both biological pathways and individual genes by calculating the gradient of the model’s predictive output with respect to the intermediate pathway layer activations and the input gene expression layer, respectively.

For a given patient sample *i*, let *ŷ*_*i*_ denote the predicted clinical outcome (risk score). We compute the gradient vectors for the gene layer 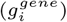 and the pathway layer 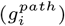 via backpropagation:

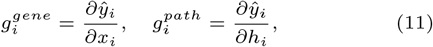

where *x*_*i*_ represents the input gene expression vector and *h*_*i*_ represents the hidden state vector of the pathway layer. The Importance Score, *S*, for a specific feature (gene *j* or pathway *k*) is defined as the absolute value of its corresponding gradient component:

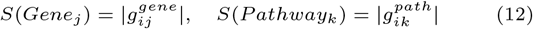

We compute these scores separately for the sex-common (*E*_*C*_) and sex-specific (*E*_*M*_, *E*_*F*_) encoders to explicitly distinguish between sex-conserved and sex-dimorphic mechanisms.

To identify robust population-level drivers, statistical testing is performed on the distribution of importance scores across the patient cohort. For each feature, a one-sample t-test is conducted to determine whether the mean importance score differs significantly from zero (*H*_0_ : *µ*_*S*_ = 0). To control for the false discovery rate (FDR) inherent in high-dimensional genomic data, *p*-values are adjusted using the Benjamini-Hochberg (BH) procedure. Features with an FDR-adjusted *p*-value < 0.01 are considered statistically significant and prioritized for further biological analysis.

## 3. Results

We evaluated FairHICON across four independent transcriptomic datasets to assess its ability to predict clinical outcomes accurately and to elucidate the underlying biological mechanisms of sexual dimorphism. We used RNA-Seq gene expression profiles to address two distinct clinical tasks: (1) cancer survival prediction and (2) asthma risk stratification. For survival prediction, we obtained data from The Cancer Genome Atlas (TCGA), focusing on Liver Hepatocellular Carcinoma (LIHC), Lung Adenocarcinoma (LUAD) and Lower Grade Brain Glioma (LGG). For risk score prediction, we utilized an asthma dataset (nasal epithelium, GSE240567) obtained from the Gene Expression Omnibus (GEO). A summary of the characteristics of the data set is provided in Table 1.

**Table 1.**
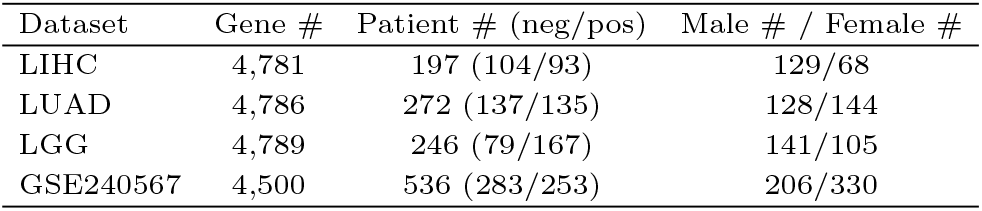
Summary of TCGA and GEO datasets used in this study.

| Dataset | Gene # | Patient # (neg/pos) | Male # / Female # |
| --- | --- | --- | --- |
| LIHC | 4,781 | 197 (104/93) | 129/68 |
| LUAD | 4,786 | 272 (137/135) | 128/144 |
| LGG | 4,789 | 246 (79/167) | 141/105 |
| GSE240567 | 4,500 | 536 (283/253) | 206/330 |

### 3.1. Comparative evaluation of predictive performance and fairness

To evaluate the predictive performance of FairHICON, we performed an extensive comparative analysis, benchmarking it against three recent methods: contrastive learning with an MLP (CL-MLP) and with XGBoost (CL-XGBoost) (Sun et al., 2024), as well as the biologically informed SPIN model (Ko et al., 2024). We divided the dataset into model training (64%), validation (16%), and testing (20%) datasets with stratified random sampling based on sex. For each experimental run, we standardized the data, scaling the validation and testing sets using the mean and standard deviation derived from the training set. We assessed performance by measuring both predictive power (i.e., Area Under the ROC Curve [AUROC] and Area Under the Precision-Recall Curve [AUPRC]) and algorithmic fairness (i.e., Demographic Parity Difference [DPD] and Equalized Odds Difference [EOD]). To ensure result stability, we replicated the experiments ten times. The quantitative results are summarized in Table 2.

**Table 2.** Comparative evaluation of predictive performance and fairness metrics across cancer (TCGA) and asthma (GSE240567) datasets. Values are reported as Mean ± Standard Deviation. **Bold** indicates the best performance, and <u>underlined</u> indicates the second-best performance. Statistical significance of FairHICON compared to the best benchmark method is denoted by asterisks: ^∗^ (*p* < 0.05) and ^∗∗^ (*p* < 0.01). Arrows indicate optimization direction (↑ higher is better; ↓ lower is better).

| Dataset | Method | Area Under ROC Curve (↑) | | | Area Under PR Curve (↑) | | | Fairness Metrics (↓) | |
| --- | --- | --- | --- | --- | --- | --- | --- | --- | --- |
|  |  | Overall | Sex-Stratified |  | Overall | Sex-Stratified |  | DPD | EOD |
|  |  |  | Male | Female |  | Male | Female |  |  |
| LGG | CL-MLP | 0.858 ± 0.040 | 0.933 ± 0.032 | 0.746 ± 0.096 | 0.844 ± 0.066 | 0.931 ± 0.037 | 0.776 ± 0.101 | 0.068 ± 0.030 | 0.206 ± 0.095 |
| | CL-XGBoost | 0.823 ± 0.066 | 0.882 ± 0.067 | 0.731 ± 0.099 | <u>0.853 ± 0.047</u> | 0.896 ± 0.058 | <u>0.810 ± 0.066</u> | **0.061 ± 0.035** | 0.230 ± 0.120 |
| | SPIN | 0.835 ± 0.062 | 0.894 ± 0.056 | 0.721 ± 0.133 | 0.811 ± 0.083 | 0.862 ± 0.076 | 0.757 ± 0.144 | 0.103 ± 0.024 | 0.283 ± 0.106 |
| | **FairHICON** | **0.892 ± 0.047\*\*** | **0.940 ± 0.044** | **0.807 ± 0.085\*\*** | **0.893 ± 0.051\*\*** | **0.938 ± 0.051** | **0.839 ± 0.082** | 0.070 ± 0.024 | **0.188 ± 0.059** |
| LIHC | CL-MLP | 0.662 ± 0.039 | 0.692 ± 0.054 | 0.581 ± 0.078 | <u>0.646 ± 0.053</u> | <u>0.703 ± 0.068</u> | 0.549 ± 0.083 | 0.085 ± 0.027 | <u>0.161 ± 0.034</u> |
| | CL-XGBoost | 0.650 ± 0.096 | 0.682 ± 0.130 | <u>0.583 ± 0.147</u> | 0.634 ± 0.098 | 0.680 ± 0.124 | 0.557 ± 0.155 | 0.121 ± 0.067 | 0.252 ± 0.122 |
| | SPIN | 0.620 ± 0.043 | 0.660 ± 0.067 | 0.526 ± 0.144 | 0.608 ± 0.065 | 0.643 ± 0.089 | <u>0.576 ± 0.118</u> | **0.076 ± 0.045** | **0.156 ± 0.098** |
| | **FairHICON** | **0.722 ± 0.056\*** | **0.762 ± 0.057** | **0.636 ± 0.140** | **0.692 ± 0.089** | **0.742 ± 0.081** | **0.634 ± 0.158** | 0.081 ± 0.043 | 0.175 ± 0.053 |
| LUAD | CL-MLP | 0.624 ± 0.051 | **0.658 ± 0.103** | 0.593 ± 0.068 | **0.628 ± 0.062** | **0.645 ± 0.123** | <u>0.636 ± 0.075</u> | **0.037 ± 0.013** | **0.077 ± 0.025** |
| | CL-XGBoost | 0.582 ± 0.053 | 0.581 ± 0.133 | 0.582 ± 0.075 | 0.585 ± 0.050 | 0.576 ± 0.139 | 0.610 ± 0.054 | 0.062 ± 0.044 | 0.116 ± 0.063 |
| | SPIN | 0.546 ± 0.043 | 0.604 ± 0.073 | 0.487 ± 0.079 | 0.541 ± 0.036 | 0.589 ± 0.097 | 0.539 ± 0.058 | <u>0.045 ± 0.040</u> | <u>0.082 ± 0.048</u> |
| | **FairHICON** | **0.638 ± 0.079** | <u>0.644 ± 0.106</u> | **0.628 ± 0.113** | <u>0.626 ± 0.079</u> | <u>0.607 ± 0.113</u> | **0.657 ± 0.107** | 0.058 ± 0.034 | 0.116 ± 0.047 |
| GSE240567 | CL-MLP | <u>0.698 ± 0.066</u> | <u>0.697 ± 0.090</u> | <u>0.699 ± 0.061</u> | 0.697 ± 0.068 | <u>0.696 ± 0.073</u> | 0.711 ± 0.079 | 0.047 ± 0.015 | 0.075 ± 0.018 |
| | CL-XGBoost | 0.692 ± 0.066 | 0.685 ± 0.093 | 0.694 ± 0.072 | 0.691 ± 0.080 | 0.693 ± 0.085 | 0.694 ± 0.097 | 0.072 ± 0.043 | 0.134 ± 0.068 |
| | SPIN | 0.681 ± 0.033 | 0.677 ± 0.059 | 0.685 ± 0.047 | <u>0.700 ± 0.041</u> | 0.689 ± 0.079 | <u>0.714 ± 0.041</u> | <u>0.038 ± 0.010</u> | <u>0.069 ± 0.015</u> |
|  | **FairHICON** | **0.734 ± 0.044\*** | **0.735 ± 0.089\*\*** | **0.734 ± 0.032** | **0.737 ± 0.052\*\*** | **0.719 ± 0.101** | **0.757 ± 0.040\*** | **0.036 ± 0.012** | **0.066 ± 0.017** |

Across all evaluated datasets, FairHICON consistently achieved the best overall predictive performance, highlighting its capacity to learn strong disease representations. In LGG dataset, our approach achieved an overall AUROC of 0.892 ± 0.047 and an AUPRC of 0.893 ± 0.051, significantly surpassing the second-best method (CL-MLP) with statistical significance (*p* < 0.01). A similar performance gain was observed on the LIHC dataset, where FairHICON achieved a notably higher AUROC than the strongest benchmark (0.722 ± 0.056 vs. 0.662 ± 0.039; *p* < 0.05). In particular, in the asthma cohort (GSE240567), FairHICON secured the highest scores on all predictive metrics, validating that the hierarchical contrastive objective captures complex genomic signals that standard baselines miss.

A primary limitation of current genomic models is the substantial performance discrepancy observed between sexes, models often perform poorly in the sex that is either less represented or biologically distinct. Table 2 reveals that benchmark methods frequently exhibit significant disparities; for instance, in the LGG dataset, CL-MLP shows a drastic performance drop between male (0.933 ± 0.032) and female (0.746 ± 0.096) cohorts. FairHICON effectively bridges this gap by specifically enhancing the representation of the underperforming group. It improved the female AUROC in LGG to 0.807 ± 0.085 (*p* < 0.01)—a marked improvement over all benchmarks—while simultaneously maintaining superior performance in the male group (0.940 ± 0.044). This capacity for unbiased prediction is most evident in the GSE240567 dataset, where FairHICON achieved near-perfect parity (Male: 0.735 ± 0.089 vs. Female: 0.734 ± 0.032), effectively eliminating the sex bias observed in competing methods.

Achieving algorithmic fairness often comes at the cost of reduced predictive accuracy (i.e., leveling down the majority group). However, FairHICON defies this trade-off by simultaneously minimizing bias metrics and maximizing utility. In terms of EOD and DPD, where lower values indicate greater fairness, our model consistently ranked among the top performers. In the LGG dataset, FairHICON significantly reduced the EOD to 0.188 ± 0.059, outperforming both the domain-specific SPIN model (0.283 ± 0.106) and CL-XGBoost (0.230 ± 0.120). Furthermore, in the asthma dataset, our model achieved the lowest bias scores across the board (DPD: 0.036 ± 0.012; EOD: 0.066 ± 0.017). These results confirm that FairHICON’s dual-objective loss function disentangles biological signals from algorithmic bias, offering a clinically viable solution for fair genomic analysis.

### 3.2. Latent space disentanglement and structure

A key advantage of FairHICON is its ability to learn representations that are simultaneously accurate and structured. This is facilitated by the two-level hierarchical contrastive objective. The first level (*h*_1_) prioritizes inter-class separability based on phenotype (clinical outcome), while the second level (*h*_2_) facilitates intra-class disentanglement based on sex attributes and representation type.

To validate the effectiveness of the representation, we visualized the latent space learned from the LGG and asthma (GSE240567) datasets (Fig. 2). The visualization shows that the model effectively separates the patient data into clear, biologically meaningful clusters that align with the outputs of the sex-common, male-specific, and female-specific encoders. This clear separation provides strong empirical evidence that the hierarchical objective, combined with dedicated encoders, effectively partitioned the learned representations according to the underlying mechanisms of sex-related diseases.

**Fig. 2.**
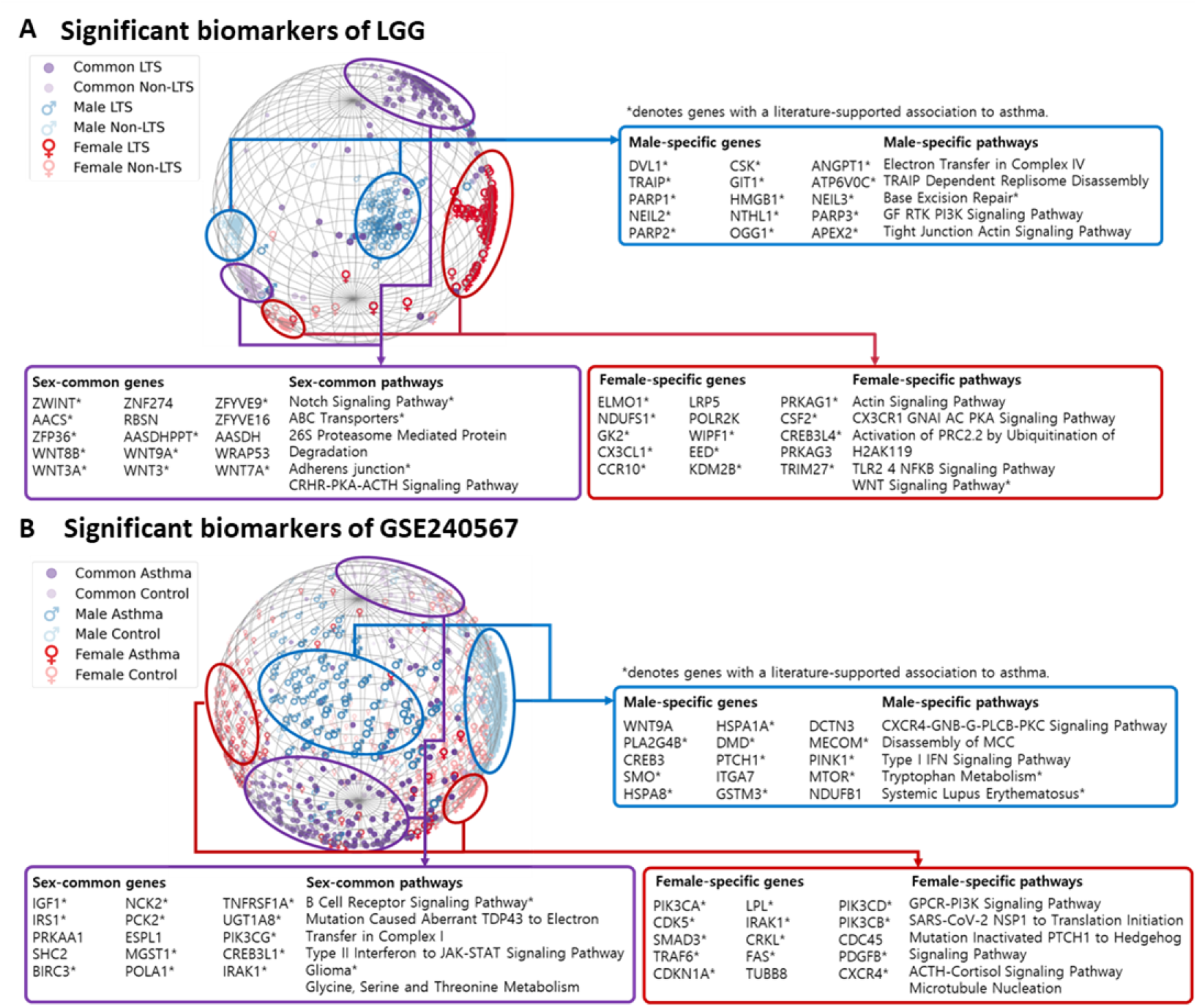
Visualization of FairHICON’s latent representations and identification of sex-specific and sex-common biomarkers. The figure displays the 3D-hypersphere latent spaces learned by FairHICON, illustrating the effective disentanglement of patient samples into distinct semantic subspaces:sex-common (purple), male-specific (blue), and memale-specific (red). (A) Analysis of the brain lower grade glioma (LGG) dataset. The spherical embedding demonstrates robust clustering of patients based on survival status (LTS vs. non-LTS), separated within each sex-specific or common subspace. The connected tables list the top-ranked genes and biological pathways identified by the model, distinguishing unique male and female risk factors from shared genomic drivers. (B) Analysis of the asthma dataset (GSE240567). The visualization shows the separation between asthma and control samples across the three sex-defined subgroups. In both panels, the tables highlight the distinct genomic mechanisms driving predictions for each group. Genes and pathways marked with an asterisk (∗) denote biomarkers with literature-supported associations to the respective disease, validating the biological relevance of the features discovered by FairHICON.

To elucidate the biological drivers underlying the model’s predictions, we performed feature attribution analysis on the representations derived from the three distinct encoders. We leveraged biologically informed architecture to identify the most influential genes and pathways and further validated these findings through statistical analysis.

### 3.3. Identification of potential significant biomarkers

FairHICON uncovered distinct groups of biomarkers that were shared between sexes and those that were sex-specific in both the LGG and GSE240567 datasets, respectively, highlighting unique genomic mechanisms underlying disease in each group. Furthermore, we verified that our findings align with known biomarkers associated with brain glioma and asthma previously reported in the biological literature, denoted by an asterisk (Fig. 2).

For the LGG analysis, we highlighted the most significant sex-common and sex-specific genes and pathways detected by FairHICON (Fig. 2A). FairHICON identified pathways involved in glioma pathogenesis regardless of sex, including *the Notch signaling pathway, ABC Transporters, 26S Proteasome Mediated Protein Degradation*, and *Adherens Junction*. The main genetic drivers associated with these shared mechanisms included *ZWINT, ZNF274, ZFYVE9, AACS*, and *WNT* family genes such as *WNT8B, WNT9A*, and *WNT3A*. In contrast, FairHICON prioritized pathways related to DNA repair and cellular energetics, such as *Electron Transfer in Complex IV, TRAIP Dependent Replisome Disassembly*, and *Base Excision Repair*. Influential genes included *DVL1, TRAIP, PARP1, NEIL2*, and *PARP2*, alongside *CSK, GIT1*, and *HMGB1*, highlighting a potential male-specific reliance on genomic stability maintenance. We revealed influential features linked to cytoskeletal organization and immune signaling. Significant pathways included the *Actin signaling pathway, CX3CR1-GNAI-AC-PKA signaling*, and *TLR2/4 NFKB signaling*. The identified key genes were *ELMO1, NDUFS1, GK2, CX3CL1*, and *CCR10*, suggesting different structural and inflammatory dependencies in the female cohort.

Similarly, for the asthma dataset (GSE240567), we present the key biomarkers uncovered by FairHICON (Fig. 2B). The shared mechanisms highlighted by the sex-common encoder involved core immune and signaling processes, such as *the B Cell Receptor signaling pathway, Type II Interferon to JAK-STAT signaling pathway*, and *Glycine, Serine and Threonine Metabolism*. The prominent genes included *IGF1, IRS1, TNFRSF1A, PCK2*, and *MGST1*, indicating conserved metabolic and inflammatory drivers. Unique to males were pathways involving *CXCR4-GNB-G-PLCB-PKC signaling, Type I IFN signaling*, and *Tryptophan Metabolism*. Top genetic drivers included *WNT9A, PLA2G4B, HSPA1A, DMD*, and *HSPA8*. The identification of *Tryptophan Metabolism* and *Type I IFN signaling* suggests specific immune-metabolic regulatory distinctness in males. The female-specific encoder emphasized pathways related to hormonal and growth factor signaling, such as *GPCR-PI3K signaling, ACTH-Cortisol signaling*, and *Microtubule Nucleation*. Key genes included *PIK3CA, CDK5, SMAD3, LPL*, and *IRAK1*. The prominence of *the ACTH-Cortisol pathway* aligns with known hormonal influences on asthma prevalence and severity in women.

### 3.4. Clinical validation of sex-specific genetic risk factors

To validate the clinical relevance of the genomic drivers identified by FairHICON (Fig.2), we evaluated their association with clinical outcomes using sex-stratified regression analyses (Fig.3). For the LGG dataset, we assessed the prognostic utility of the prioritized genes via Cox Proportional Hazards (CPH) regression (Fig.3A); for the asthma dataset, we evaluated disease susceptibility using logistic regression to determine Odds Ratios (OR) (Fig.3B).

**Fig. 3.**
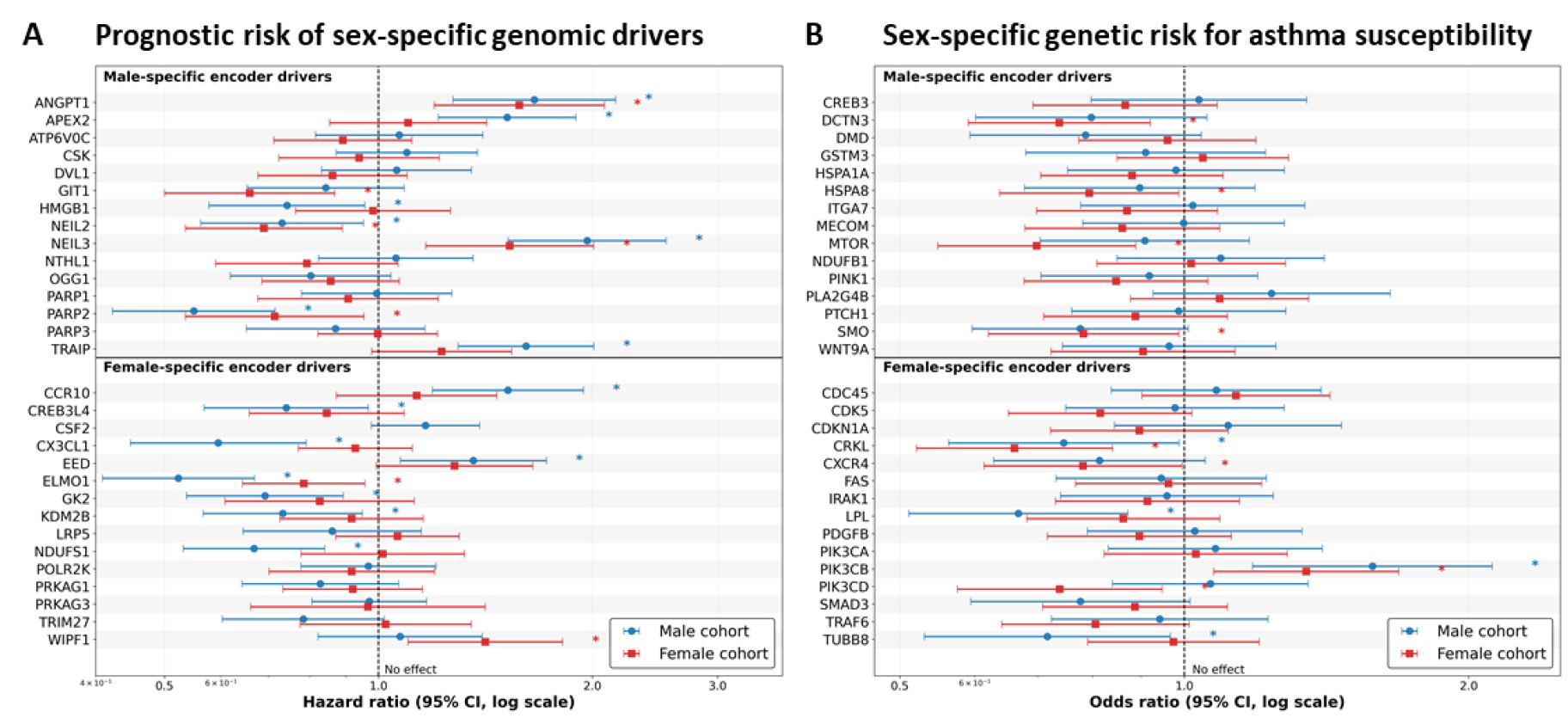
Quantitative assessment of sex-specific genomic drivers identified by FairHICON. (A) Prognostic risk analysis for LGG. Forest plots display the Hazard Ratios (HR) with 95% confidence intervals (CI) for top genes identified by the Male-Specific Encoder (top panel) and Female-Specific Encoder (bottom panel). Risks are stratified by sex (Blue: Male Cohort, Red: Female Cohort). HR *>* 1 indicates increased risk of mortality, while HR < 1 indicates a protective effect. (B) Genetic risk analysis for asthma susceptibility (GSE240567). Forest plots display the Odds Ratios (OR) with 95% CI for top genes, stratified by sex. OR *>* 1 indicates increased susceptibility to asthma. The vertical dashed line represents the no effect threshold (HR/OR = 1). An asterisk (∗) marks results that are statistically significant at *p* < 0.05.

For the prognosis of LGG cancer, the analysis revealed a substantial dissociation in risk attribution between the sexes. First, genomic factors prioritized by the male-specific encoder revealed distinct risk profiles. *NEIL3* and *ANGPT1* emerged as robust risk factors, exhibiting Hazard Ratios (HR) *>* 1 in both sexes, with *NEIL3* showing a particularly strong association in males (HR ≈ 2.0, *p* < 0.05). *TRAIP* demonstrated a distinct sex-specific effect, functioning as a significant risk factor for males (HR *>* 1, *p* < 0.05) while showing no statistically significant association in females (95% CI crossing 1.0). Conversely, genes such as *PARP2, PARP3*, and *OGG1* displayed protective associations (HR < 1) in the male cohort, indicating their higher expression correlates with better survival outcomes. Second, we found that the identified female-specific drivers exerted particularly strong effects in women. *WIPF1* was a standout female-specific risk factor, exhibiting a significantly elevated HR in the female cohort compared to a negligible effect in males (HR *>* 1, *p* < 0.05). *ELMO1* and *NDUFS1* were identified as robust protective factors (HR < 1) for both sexes, with non-overlapping confidence intervals suggesting a consistent beneficial prognostic value.

Extending this validation to disease susceptibility in asthma (GSE240567), FairHICON identified several genes associated with increased asthma risk in males, including *PLA2G4B* and *NDUFB1*, both showing OR *>* 1. *GSTM3* and *DMD* exhibited protective effects (OR < 1) within the male cohort, suggesting that their expression may be associated with reduced susceptibility. Genes identified through the female encoder were strongly linked to asthma susceptibility. *PIK3CB* displayed a remarkably high odds ratio in the female cohort, indicating that it is a critical susceptibility factor for women (OR *>* 1, *p* < 0.05). In contrast, *PIK3CD* and *CXCR4* were associated with a reduced susceptibility (OR < 1, *p* < 0.05) in the female cohort, highlighting their potential protective roles in the pathology of female asthma.

### 3.5. Validation of sexual dimorphism via prognostic stratification

To empirically the sexual dimorphism identified by FairHICON, we evaluated the prognostic distinctiveness of the prioritized pathways. We hypothesized that if the model disentangles sex-specific pathology, a pathway identified as a male-specific driver should stratify survival outcomes significantly better in the male cohort than in the female cohort, while pathways identified as female-specific should demonstrate the inverse pattern. Kaplan-Meier (KM) survival analysis on the LGG dataset confirmed this hypothesis (Fig. 4).

**Fig. 4.**
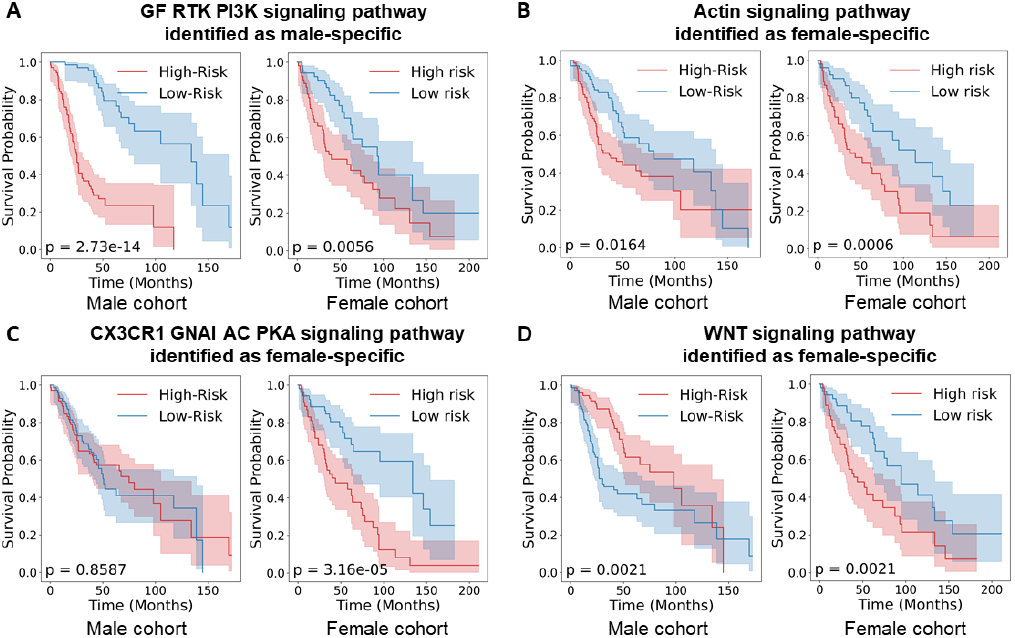
Validation of sexual dimorphism through comparative survival analysis. Patients were stratified into high-risk (red) and low-risk (blue) groups based on the median activation of sex-specific pathways. The plots demonstrate that pathways stratify survival more effectively in the sex for which they were identified. (A) Male-specific driver: *GF RTK PI3K signaling pathway* exhibits a profound risk separation in the male cohort (*p* = 2.73 × 10^−14^), significantly outperforming the stratification observed in the female cohort (*p* = 0.0056). (B-D) Female-specific driver: *Actin signaling pathway* differentiates risk more significantly in females (*p* = 0.0006) than in males (*p* = 0.0164). *CX3CR1 GNAI AC PKA signaling pathway* functions as a female-exclusive prognostic marker (*p* < 10^−4^), showing no stratification capability in the male cohort (*p* = 0.8587). *WNT signaling pathway* similarly displays superior prognostic stratification in the female cohort relative to the male cohort. *p*-values were calculated using the log-rank test.

The *GF RTK PI3K signaling pathway* was identified as a significant risk factor by the male-specific encoder. Consistent with this attribution, the pathway exhibited exceptional prognostic power within the male cohort, yielding a clear separation between high-risk and low-risk groups (*p* = 2.73 × 10^−14^). In contrast, while the pathway remained statistically associated with survival in females (*p* = 0.0056), the stratification was orders of magnitude less distinct than in males. This disparity indicates that while this signaling axis is relevant to glioma generally, its specific dysregulation constitutes a disproportionately critical survival determinant for male physiology. This computational finding aligns with established experimental evidence indicating that male glioblastoma is disproportionately driven by cell cycle and growth factor signaling. Specifically, male glioblastoma cells exhibit significantly higher activity in the PI3K/mTOR pathway compared to female cells (Sponagel et al., 2022). Furthermore, glycolytic signatures downstream of this pathway yield significant prognostic value in males while failing to stratify survival in females (Ippolito et al., 2017), providing robust external validation for the male-specific risk attribution identified by FairHICON.

In contrast, pathways prioritized by the female-specific encoder, such as *Actin signaling, CX3CR1 GNAI AC PKA signaling*, and *WNT signaling*, demonstrated superior prognostic utility in women. The most definitive evidence of dimorphism was observed in the *CX3CR1 GNAI AC PKA signaling pathway*. This mechanism successfully stratified the female cohort with high significance (*p* < 10^−4^) but failed entirely to predict survival outcomes in the male cohort (*p* = 0.8587). Similarly, the *Actin signaling pathway* achieved a highly significant separation in females (*p* = 0.0006) compared to a weaker association in males. This mirrors the findings that the survival of female glioblastoma is more closely related to invasive and structural signaling profiles (such as integrin and actin dynamics) than the predominant cell-cycle drivers in males (Yang et al., 2019).

These comparative survival analyzes confirm that the features extracted by FairHICON are not generic cancer markers, but rather distinct biological drivers that reflect the underlying sexual dimorphism of the disease. By correctly assigning the PI3K axis to males and structural/invasive pathways to females, the model demonstrates true mechanistic validity, ensuring that clinical risk stratification is tailored to the distinct biological reality of each sex.

## 4. Discussion

Sex-based differences are crucial for understanding how complex human disease incidence, progress, and treatment response, yet it remains an underrepresented factor in the design of AI-driven predictive models. The tendency of AI models to exhibit algorithmic bias, particularly when trained on imbalanced genomic datasets, poses a significant challenge to the fair application of these technologies in oncology. In this study, we introduced FairHICON, a novel approach specifically designed to address fairness in the context of sexual dimorphism analysis using gene expression data. By integrating supervised hierarchical contrastive learning with biologically informed architectures, FairHICON advances the dual objectives of enhancing predictive accuracy and mitigating bias.

The core innovation of FairHICON lies in its two-level hierarchical contrastive learning strategy. FairHICON extends supervised contrastive learning by introducing a secondary level of optimization (*h*_2_) that uses group-normalized loss (GN-SupCon) to ensure fairness between sex subgroups within each class. By first separating representations based on clinical outcomes (*h*_1_) and then addressing intra-class variation equitably (*h*_2_), the approach effectively prevents prioritization of the majority sex group, a common pitfall in standard machine learning approaches. This structured learning process ensures that the resulting representations are highly predictive and unbiased. Our empirical evaluation across four diverse datasets demonstrates the superiority of FairHICON. FairHICON consistently achieved the highest overall AUC while simultaneously maintaining excellent fairness metrics (DPD and EOD). The analysis of subgroup performance revealed that FairHICON enhanced predictive performance for both male and female cohorts in most datasets. This confirms that the integration of fairness constraints within the FairHICON approach does not necessitate a trade-off in predictive power; rather, it leads to more robust and generalizable models by learning less biased representations.

The interpretability of AI models is paramount for their translation into clinical practice. FairHICON addresses this need through its biologically informed architecture (BINN) and the use of a Siamese structure with distinct encoders. The incorporation of pathway knowledge constrains the learning process to biologically plausible mechanisms. The distinct encoders, combined with robust attribution methods, allow for the direct identification and comparison of sex-common and sex-specific biological risk factors. As demonstrated in the analysis of the GSE240567 and LGG datasets, FairHICON disentangled the latent space into meaningful clusters, driven by genes and pathways with known relevance, such as distinct immune and hormonal signaling pathways in males and females. This capability moves beyond black-box predictions, offering actionable biological insights into the molecular mechanisms underlying sexual dimorphism.

Despite these promising results, several limitations warrant consideration. The analysis of bulk gene expression data inherently involves the High-Dimensional Low-Sample Size (HDLSS) challenge, which can affect the robustness and generalizability of the findings. Although we employed preprocessing strategies and pathway integration to mitigate overfitting, this risk remains an inherent challenge in genomic studies. Furthermore, reliance on predefined pathway databases can limit the discovery of novel biological connections outside existing knowledge graphs. Future work will aim to address these constraints by applying FairHICON to single-cell RNA-Seq data, providing higher-resolution insights into cellular heterogeneity. Furthermore, extending the framework to integrate multi-omics data, including genomics, epigenomics, and proteomics, provide a more comprehensive understanding of the molecular landscape of sexual dimorphism in cancer.

## 5 Acknowledgments

This research was supported by the National Science Foundation Major Research Instrumentation (NSF MRI) (Grant#: 2117941).

## Notes

### Competing Interest Statement

The authors have declared no competing interest.

